# Stronger net selection on males across animals

**DOI:** 10.1101/2021.04.16.440171

**Authors:** Lennart Winkler, Maria Moiron, Edward H. Morrow, Tim Janicke

## Abstract

Sexual selection is considered the major driver for the evolution of manifold sex differences. However, the eco-evolutionary dynamics of sexual selection and their role for a population’s adaptive potential to respond to environmental change have only recently been explored. Theory predicts that sexual selection promotes adaptation at a low demographic cost only if net selection is stronger on males compared to females. We used a comparative approach to show that net selection is indeed stronger in males in species prone to intense sexual selection. Given that both sexes share the vast majority of their genes, our findings corroborate the notion that the genome is often confronted with a more stressful environment when expressed in males. Collectively, our study supports a long-standing key assumption required for sexual selection to bolster adaptation, and intense sexual selection may therefore enable some species to track environmental change more efficiently.

**One sentence summary:** Comparative study finds support for stronger net selection in males.

## Introduction

For almost a century researchers have gathered compelling evidence that sexual selection (i.e. selection arising from competition for mating partners and/or their gametes) constitutes the ultimate evolutionary force generating sexual dimorphism in a multitude of reproductive characters and life-history traits (Andersson 1994; Clutton-Brock 2007). Despite this outstanding progress, we are just beginning to understand the eco-evolutionary dynamics of sexual selection in terms of its impact on demography and adaptive potential of a population (Svensson 2019). Traditionally, sexual selection has often been considered to impair population growth and constrain the adaptation to changing environments as a consequence of inter- and intra-locus sexual conflict (Arnqvist & Rowe 2005). More recently, however, both theoretical and empirical work suggest that sexual selection can facilitate how populations cope with environmental change (Lorch *et al*. 2003; Candolin & Heuschele 2008; Holman & Kokko 2013; Martínez-Ruiz & Knell 2017; Martinossi-Allibert *et al*. 2019). This latter school of thought is based on two main assumptions. First, sexual and natural selection need to be aligned, meaning that sexual selection favors alleles that also improve fecundity and survival − a process mediated by condition-dependence of sexually selected traits (Rowe & Houle 1996). Such an alignment is expected to manifest in a positive genetic correlation between male and female fitness components, which has already been documented across animal taxa (Poissant *et al*. 2010), though there is also evidence for negative cross-sex genetic correlations indicating the presence of intra-locus sexual conflict in some species (Chippindale *et al*. 2001). Moreover, there is also solid comparative and meta-analytic evidence supporting that the expression of pre− and post−copulatory sexual traits depends on the overall condition of the male (Cotton *et al*. 2004; Macartney *et al*. 2019) implying that sexual selection may not only favor the evolution of prominent secondary sexual traits but also traits that confer health and vigor (Jennions *et al*. 2001), and therefore eventually purges deleterious alleles.

In contrast to the solid support for an alignment of sexual and natural selection in many species, we still know very little on whether the second key assumption for sexual selection to empower evolutionary adaptation is generally fulfilled across a broad range of taxa. That is that sexual selection enforces natural selection only if it gives rise to stronger net selection (defined as the sum of genome-wide selection against deleterious alleles) on males compared to females. In a landmark synthesis paper, Whitlock and Agrawal (2009) explore the effect of sex-specific selection on the population’s mutation load, i.e. the overall reduction of absolute fitness due to deleterious alleles in a population. They expanded the fundamental work of Haldane (1937) by relaxing the assumption of random mating to demonstrate that the mutation load *L* is expected to be

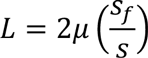

where *μ* is the mutation rate from good to bad alleles and *s* is the average selection coefficient against deleterious alleles of females (*s_f_*) and males. Hence, the mutation load is reduced whenever *s_f_* < *s*, which arises if net selection is stronger on males compared to females. Given that the population’s productivity is typically governed by female fecundity, a population with stronger net selection on males can purge its mutation load and adapt faster to a new environment with a lowered demographic cost and thereby reducing its extinction risk. In other words, females benefit from being part of a gene pool that is purified primarily through stronger selection on males (Whitlock & Agrawal 2009).

There is ample empirical evidence that typically (though not always) males undergo stronger sexual selection whereas females are primarily exposed to fecundity selection (Janicke *et al*. 2016) as predicted by Bateman’s principle (Bateman 1948). However, our knowledge on whether such stronger sexual selection on males eventually translates into stronger net selection relative to females is still limited and equivocal (Whitlock & Agrawal 2009; Hendry *et al*. 2018). This lack of evidence stems at least partially from difficulties in quantifying the strength of net selection in a framework that also allows comparisons among sexes and species. Potentially the most promising metric to contrast the strength of net selection across contexts is the mean-standardized variance in fitness, which is often expressed as the coefficient of variation (*CV*). Previous comparative studies provided evidence that the phenotypic variance in fitness (‘opportunity for selection’ (Crow 1958)) is typically larger in males compared to females, suggesting that the upper opportunity for net selection is stronger in males (Janicke *et al*. 2016). However, it has been questioned whether the phenotypic variance in fitness provides a good proxy for the strength of net selection because environmental variation can substantially inflate this variance, which limits its explanatory power with respect to evolutionary responses and complicates the comparison across contexts and studies (Whitlock & Agrawal 2009). To overcome this problem, researchers have advocated to use the genetic rather than the phenotypic variance in fitness as a proxy for the strength of net selection (Jones 2009; Hendry *et al*. 2018). This is also because the genetic variance in relative fitness corresponds to the rate of increase in mean fitness that results from selection on allele frequencies (Fisher 1930). Therefore, the mean-standardized genetic variance (*CV*_G_) in fitness provides a highly diagnostic metric to quantify the strength of net selection, but it has to our knowledge not yet been subjected to a systematic and global test for an overall sex difference.

Here we addressed this key aspect of sexual selection theory by using a comparative approach to test whether net selection is generally stronger on males across a broad taxonomic range. Specifically, we ran a systematic literature search and compiled 101 pairwise estimates of male and female genetic variances for two main fitness components – reproductive success and lifespan – from a total of 26 species. Applying phylogenetically-informed comparative analyses we tested (*i*) whether phenotypic variances are aligned to genetic variances as assumed by the phenotypic gambit (Grafen 1991) and (*ii*) whether genetic variances show consistent sex differences with the prediction that males show larger genetic variance in reproductive success but not lifespan. According to the so-called *genic capture hypothesis,* virtually every deleterious allele impairs the condition of an organism, which is defined as the pool of acquired resources that can be allocated into different fitness components such as survival and reproductive success (Rowe & Houle 1996). As a consequence, all fitness components are expected to be condition-dependent, i.e. to positively covary with condition because it captures the overall genetic quality of a given organism. Importantly, if sexual selection operates on males, reproductive success in males does not only depends on gamete production and post-zygotic investment (as predicted to constitute the primary determinants in females), but also on the outcome of pre- and post-copulatory mate choice and mate competition. Therefore, a deleterious allele with its negative effect on condition is predicted to have a disproportional higher impact on reproductive success in males compared to females. This heightened condition dependence of male performance (Rowe & Houle 1996) is expected to translate ultimately into a higher genetic variance in reproductive success in males (Whitlock & Agrawal 2009; Hendry *et al*. 2018). By contrast, lifespan is considered to be unaffected by the outcome of sexual selection except in the relatively rare event of mortal combats for access to mates, mate harassment or indirectly by shaping life-history strategies (Bonduriansky *et al*. 2008). If correct, a deleterious allele is therefore expected to have a similar effect on lifespan in males and females so that we predict if anything, a much weaker male bias for genetic variance in lifespan. In this context, we further explored the role of sexual selection for generating the hypothesized sex differences in net selection. For this purpose we contrasted socially monogamous and polygamous species, with the prediction that a male bias in genetic variance for reproductive success is primarily prevalent in polygamous species where sexual selection is most likely to be stronger compared to monogamous species (Shuster & Wade 2003).

## Material and Methods

### Literature Search

We ran a systematic literature search in order to obtain an unbiased sample of coefficients of genetic variation for two major fitness components: reproductive success and lifespan. Specifically, we screened the ISI Web of Science Core Collection database (Clarivate Analytics) on 2^nd^ of August 2019 (for search terms see Supplementary Material). This search yielded 3793 records. Moreover, we screened previous synthesis articles on related questions (Poissant *et al*. 2010; Hendry *et al*. 2018; Connallon & Matthews 2019) for other primary studies by which we could identify one additional record (Wheelwright *et al*. 2014). Furthermore, we posted a request on ‘evoldir’ mailing list (http://life.mcmaster.ca/evoldir.html) and ResearchGate platform (https://www.researchgate.net/) for unpublished data, which resulted in one additional study (Abbott and Norden, in prep.). Finally, we added two published studies indicated by colleagues (Gay *et al*. 2011; Pélissié *et al*. 2012) and two of our own unpublished studies (Janicke et al., in prep.; Moiron et al., in prep.). After exclusion of duplicates and screening of abstracts, we checked a total of 203 records for eligibility based on three selection criteria. First, studies must report or include information to compute coefficients of genetic variation of reproductive success and/or lifespan. Second, studies must report genetic parameters for males and females both quantified under same conditions (i.e. same field populations or same experimental laboratory conditions). Third, we only included studies on animals simply because of the scarcity of data on genetic variances of male and female fitness components in plants. The final dataset included data from 55 primary studies (see Supporting Information for PRISMA diagram (Fig. S1) and reference list of all primary studies).

We note that primary studies differed in terms of how genetic variances were estimated including full-sib breeding designs (*N* = 3), half-sib breeding designs (*N* = 12), inbred lines (*N* = 17), pedigrees (*N* = 21) and twin studies (*N* = 2). While this may have induced potential biases in absolute values, we do not expect that the different experimental approaches led to a systematic bias in the sex difference of genetic variance. Therefore, we pooled all data obtained from different approaches in all analyses.

### Data acquisition

For all primary studies we extracted four parameters for both sexes: (*i*) sample size, (*ii*) arithmetic mean, (*iii*) phenotypic variance, and (*iv*) genetic variance of reproductive success and/or lifespan. For 11 studies at least one of these parameters was not reported in the article. In these cases, we received the parameter estimates from the authors upon request or reanalyzed the raw data (either published together with the article or provided by the authors). We computed the coefficients of phenotypic and genetic variation (*CV*_P_ and *CV*_G_, respectively) as the square root of the variance (i.e. the standard deviation) divided by the arithmetic mean, which makes this metric comparable across contexts and species. Note that *CV* of a given trait is often denoted as ‘evolvability’ (Houle 1992) and equals the square root of the opportunity for selection *I*, which is also frequently used to quantify the upper limit of the strength of selection (Jones 2009; Hendry *et al*. 2018).

In total we obtained 101 paired estimates of *CV*_P_ and *CV*_G_ for males and females, including 62 estimates for reproductive success and 39 estimates for lifespan. In all primary studies, reproductive success was measured as the number of offspring except for one study in which it was estimated as the number of grandchildren (Bolund *et al*. 2013). Lifespan was primarily measured as adult survival (i.e. excluding mortality until reaching maturity; 33 out of 39 estimates) and in the few remaining cases as the age of last reproduction, reproductive lifespan (calculated as time between first and last reproduction), or total lifespan (including juvenile mortality).

### Phylogeny and Mating System Classification

The 55 primary studies encompass a total of 26 animal species with an overrepresentation of insects (*N* = 12) and birds (*N* = 7). In order to account for any source of phylogenetic non-independence we reconstructed the phylogeny of all sampled species (Fig. S2). Firstly, we retrieved pairwise estimates of divergence times from the TimeTree database (http://www.timetree.org/; (Kumar *et al*. 2017)). Secondly, we aged undated nodes on the basis of divergence times of neighboring nodes applying the branch length adjuster (BLADJ) algorithm (Webb *et al*. 2008). Finally, we used the resulting distance matrix to compute a phylogeny, using the unweighted pair group method with arithmetic mean (UPGMA) algorithm implemented in MEGA (https://www.megasoftware.net/; (Kumar *et al*. 2018)) and transformed it into the Newick format for further analysis.

To explore the role of sexual selection in generating sex differences in phenotypic and genetic variances, we used published information in the scientific literature on the social mating system as a proxy for the strength of sexual selection. Specifically, we distinguished between socially monogamous (*N* = 6) and polygamous species (*N* = 20; including polygynous and polygynandrous species). In strictly monogamous species the variance in reproductive success is expected to be identical for males and females. However strict monogamy is rather rare in animals (Lukas & Clutton-Brock 2013) and most of our sampled monogamous species are birds, which are described as socially monogamous rather than genetically monogamous because all of them show at least some degree of extra-pair paternity (Brouwer & Griffith 2019). Moreover, the occurrence of partial polygyny in some sampled bird species rendered their classification problematic. In these problematic cases, we searched the literature for estimates of the proportion of polygynous males in the studied population and only considered those species with less than 10 % of polygynous males as socially monogamous (Table S1). Hence, even though sexual selection is likely to operate also in most socially monogamous species, we assume that the strength of pre- and post-copulatory sexual selection is generally higher in polygamous compared to monogamous species (Shuster & Wade 2003). We examined the sensitivity of our classification criteria to different thresholds that have been previously used by other authors and found that the identity of monogamous and polygamous species remained when applying a 5% (Moller 1986) or a 15% threshold (Dunn *et al*. 2001). Nevertheless, we acknowledge that our classification is an oversimplification of a continuum in the strength of sexual selection across species in nature and we stress that all provided tests for correlations with the social mating system are exploratory, especially given the underrepresentation of monogamous species and their taxonomically uneven distribution in our dataset.

**Table 1.** Results of Phylogenetic General Linear Mixed Effect Models testing for an effect of sex on phenotypic (*CV*_P_) and genetic (*CV*_G_) coefficient of variation. Results shown for reproductive success (RS) and lifespan (LS) for models ran across mating systems and when ran separately for socially monogamous and polygamous species. Estimates are shown as posterior means with 95% Highest Posterior Density (HPD) intervals, with positive values indicating a male bias. The variance explained by sex is given as the marginal *R*^2^ and the phylogenetic signal is reported as *H*^2^.

| Response | Variance component | Sex effect estimate | $P_{MCMC}$ | Marginal $R^2$ | Phylogenetic $H^2$ |
| --- | --- | --- | --- | --- | --- |
| Across mating systems |  |  |  |  |  |
| RS | $CV_P$ | 0.234 (0.149, 0.322) | < 0.001 | 0.07 (0.02, 0.12) | 0.09 (0.00, 0.28) |
| | $CV_G$ | 0.087 (0.043, 0.129) | < 0.001 | 0.04 (0.01, 0.09) | 0.21 (0.01, 0.53) |
| LS | $CV_P$ | 0.002 (-0.024, 0.027) | 0.885 | < 0.01 (0.00, 0.00) | 0.37 (0.02, 0.74) |
| | $CV_G$ | 0.019 (-0.002, 0.042) | 0.094 | 0.01 (0.00, 0.02) | 0.37 (0.03, 0.77) |
| Monogamy |  |  |  |  |  |
| RS | $CV_P$ | -0.012 (-0.057, 0.034) | 0.580 | < 0.01 (0.00, 0.00) | 0.29 (0.00, 0.95) |
| | $CV_G$ | -0.015 (-0.076, 0.047) | 0.609 | 0.01 (0.00, 0.03) | 0.33 (0.01, 0.90) |
| LS | $CV_P$ | 0.027 (-0.014, 0.068) | 0.170 | < 0.01 (0.00, 0.02) | 0.27 (0.00, 0.83) |
| | $CV_G$ | 0.050 (-0.021, 0.123) | 0.148 | 0.03 (0.00, 0.10) | 0.43 (0.01, 0.87) |
| Polygamy |  |  |  |  |  |
| RS | $CV_P$ | 0.313 (0.209, 0.417) | < 0.001 | 0.11 (0.04, 0.19) | 0.17 (0.00, 0.45) |
| | $CV_G$ | 0.119 (0.067, 0.170) | < 0.001 | 0.07 (0.02, 0.14) | 0.17 (0.00, 0.50) |
| LS | $CV_P$ | -0.006 (-0.036, 0.028) | 0.718 | < 0.01 (0.00, 0.00) | 0.61 (0.20, 0.94) |
| | $CV_G$ | 0.009 (-0.013, 0.031) | 0.390 | 0.01 (0.00, 0.02) | 0.33 (0.03, 0.71) |

### Statistical Analyses

Statistical analyses were carried out in two steps. First, we examined the key assumption of the ‘phenotypic gambit’ by testing whether estimates of phenotypic variance predict the estimated genetic variance. For this we computed the Pearson correlation coefficient *r*, testing the relationship between *CV*_P_ and *CV*_G_ for both sexes and the two fitness components separately. In addition, we tested whether the sex bias in *CV*_P_ translates into a sex bias in *CV*_G_ by correlating the coefficient of variation ratio lnCVR (Nakagawa *et al*. 2015), which refers to the ln-transformed ratio of male *CV* to female *CV*, with positive values indicating a male bias. The analyses on the phenotypic gambit were motivated from a methodological perspective and we did not expect that inter-specific variation in the difference between *CV*_P_ and *CV*_G_ can be explained by a shared phylogenetic history. However, for completeness, we also ran correlations on phylogenetic independent contrasts (PICs; computed using the *ape* R-package (version 5.4.1) in R (Paradis & Schliep 2019)) to test whether our findings were robust when accounting for potential phylogenetic non-independence. We report Pearson’s correlation coefficients *r* for normally distributed data and Spearman’s *ρ* if assumptions of normality were violated.

Second, we tested the hypothesis that net selection is stronger on males by testing for a male bias in *CV*_P_ and *CV*_G_. Specifically, we ran Phylogenetic General Linear Mixed-Effects Models (PGLMMs) with *CV*_P_ or *CV*_G_ as the response variable, and sex as a fixed effect. To account for the paired data structure, we added an observation identifier as a random effect. Moreover, all models included a study identifier and the phylogeny (transformed into a correlation matrix) as random effects to account for statistical non-independence arising from shared study design or phylogenetic history, respectively. Note that the latter also accounts for the non-independence of estimates obtained from the same species as some studies estimated genetic variances from distinct field populations (Fox *et al*. 2004) or different experimental treatments under laboratory conditions such as food stress (Holman & Jacomb 2017) and temperature stress (Berger *et al*. 2014). In an additional series of PGLMMs we tested whether our proxy of sexual selection explained inter-specific variation in the sex-differences of *CV*_P_ or *CV*_G_ by adding mating system and its interaction with sex as fixed effects to the models. Finally, given that primary studies varied in the empirical approach used to quantify *CV*_P_ and *CV*_G_, we also used PGLMMs to test whether study type (23 field studies *versus* 32 laboratory studies) represented a methodological determinant of the observed sex-differences in *CV*_P_ and *CV*_G_.

All PGLMMs were ran with the *MCMCglmm* R-package (version 2.2.9) (Hadfield 2010), using uninformative priors (*V* = 1, *nu* = 0.002) and an effective sample size of 20000 (number of iterations = 11000000, burn-in = 1000000, thinning interval = 500). We computed the proportion of variance explained by fixed factors (‘marginal *R*^2^’) (Nakagawa & Schielzeth 2013). In addition, we quantified the phylogenetic signal as the phylogenetic heritability *H*^2^ (i.e., proportional variance in *CV*_P_ or *CV*_G_ explained by species identity), which is equivalent to Pagel’s ((de Villemereuil & Nakagawa 2014).

In a previous study testing for sex-specific phenotypic variances in reproductive success (Janicke *et al*. 2016), we ran formal meta-analyses using lnCVR as the tested effect size (Nakagawa *et al*. 2015). This is potentially a more powerful approach for comparing phenotypic variances but rendered unsuitable when comparing genetic variances. This is because the computation of the sampling variance of lnCVR is a function of the sample size of the sampled population and the point estimate of lnCVR (Nakagawa *et al*. 2015). However, genetic variances are estimates from statistical models and notorious for being estimated with low precision (i.e. have large confidence intervals). Therefore, using a meta-analytic approach for genetic variances using lnCVR as an effect size leads to overconfident estimation of the global effect size and is therefore likely to result in type-II-errors. However, to allow comparison with the previous meta-analysis, we report the outcome of phylogenetic meta-analyses on phenotypic variances using lnCVR in the Supplementary Material (Table S2), which largely reflects the results on the point estimates of *CV*_P_ from PGLMMs.

## Results

We found that the phenotypic coefficient of variation (*CV*_P_) of reproductive success does not predict the genetic coefficient of variation (*CV*_G_) in either males (raw estimate: *r* = 0.18, *N* = 62, *P* = 0.160; Phylogenetic Independent Contrasts (PICs): *r* = 0.20, *N* = 61, *P* = 0.126) or females (raw estimate: *r* = 0.08, *N* = 62, *P* = 0.557; PICs: *r* = 0.06, *N* = 61, *P* = 0.623; Fig. S3). In contrast, we detected a significant positive correlation between *CV*_P_ and *CV*_G_ for lifespan in males (raw estimate: *r* = 0.45, *N* = 39, *P* = 0.004; PICs: *ρ* = 0.60, *N* = 38, *P* < 0.001) and females (raw estimate: *r* = 0.45, *N* = 39, *P* = 0.005; PICs: *ρ* = 0.40, *N* = 38, *P* = 0.014). Despite these distinct findings for both fitness components, we found that the sex bias in *CV*_P_ and *CV*_G_ quantified as lnCVR was positively correlated for reproductive success (raw estimate: *r* = 0.41, *N* = 62, *P* < 0.001; PICs: *ρ* = 0.47, *N* = 61, *P* < 0.001) and lifespan (raw estimate: *r* = 0.37, *N* = 39, *P* = 0.020; PICs: *ρ* = 0.34, *N* = 38, *P* = 0.039).

Most importantly, *CV*_P_ of reproductive success was generally larger in males compared to females, which translated into a male bias in *CV*_G_, with sex explaining 6 % and 4 % of the observed variance, respectively (Table 1; Fig. 1A-B and Fig. 2). Remarkably, this sex difference could be detected in polygamous but not monogamous species, which manifested in a significant sex by mating system interaction (Table 1; Fig. 1 and Fig. 2). Contrary to the results for reproductive success, we did not observe consistent sex differences in *CV*_P_ and *CV*_G_ for lifespan (Table 1; Fig. 1C-D2). Finally, study type did not predict the sex difference in *CV*_P_ and *CV*_G_ neither for reproductive success nor lifespan (Table S3).

**Fig. 1.**
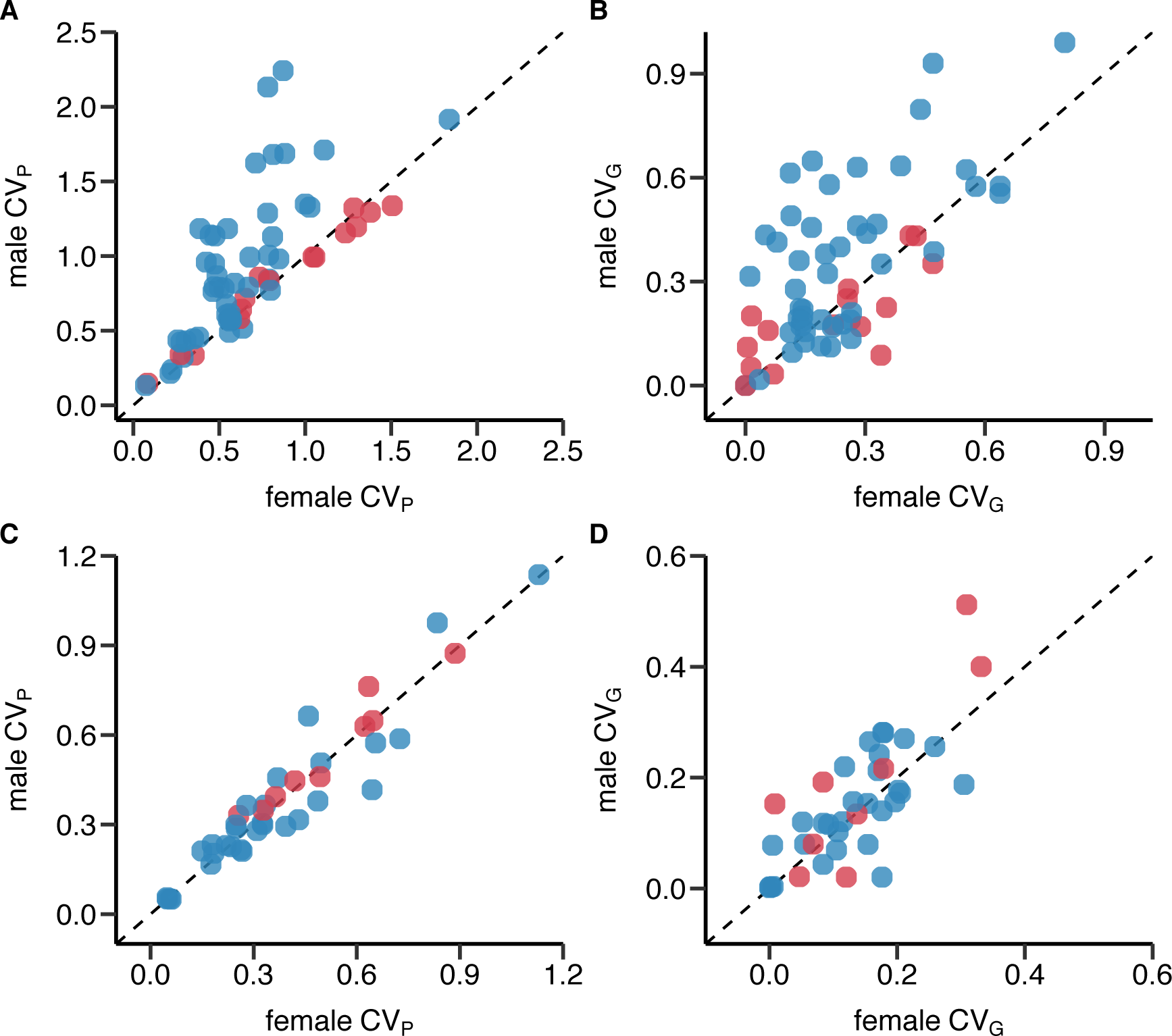
Male bias in phenotypic and genetic variances in reproductive success but not lifespan. Scatterplots show the coefficient of phenotypic variation *CV*_P_ (A, C) and genetic variation *CV*_G_ (B, D) for reproductive success (A, B) and lifespan (C, D). Monogamous species are represented in red, polygamous species in blue. All data points above the diagonals indicate a male bias.

**Fig. 2.**
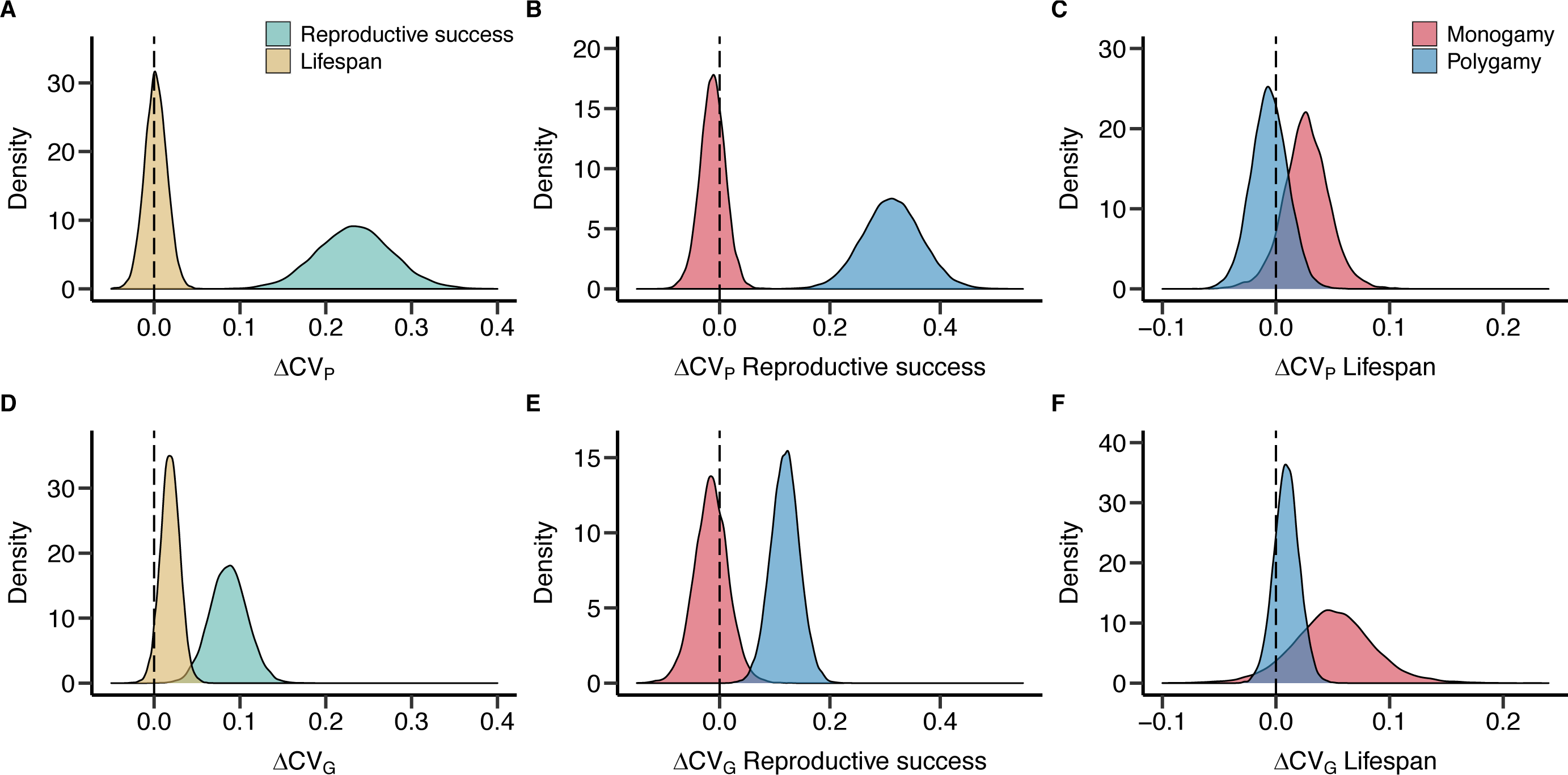
Sex differences in phenotypic and genetic coefficients of variation for reproductive success and lifespan. Plots show posterior distributions for the sex difference of the phenotypic (A-C) and genetic (D-F) coefficient of variation (Δ*CV*_P_ and Δ*CV*_G_, respectively) obtained from PGLMMs (see Methods section). Positive values indicate a male, negative values a female bias. Density plots contrast fitness components pooled across mating systems (A and D) or compare socially monogamous and polygamous species separately for reproductive success (B and E) and lifespan (C and F).

## Discussion

Males and females share the vast majority of their genome but are often subject to fundamentally different selection pressures, which is predicted to impact the demography and the adaptive potential of a population when facing environmental change (Whitlock & Agrawal 2009; Holman & Kokko 2013; Svensson 2019). In line with sexual selection theory, our study provides first comparative evidence that genome-wide selection is generally stronger on males compared to females. More specifically, our results have two major implications. First, phenotypic variance of lifespan, but not reproductive success, is aligned to genetic variance, suggesting that the phenotypic gambit does not hold for the latter. Therefore, the opportunity for selection measured as the phenotypic variance in reproductive success is a poor predictor for the strength of net selection. Despite this, the sex difference in phenotypic variance is positively correlated with the sex difference in genetic variance in both fitness components. As a consequence, our second major finding is that the previously observed male bias in the phenotypic opportunity for selection (Janicke *et al*. 2016) is also reflected in an overall higher male genetic variance in reproductive success. Importantly, this male bias can only be detected in polygamous species, in which sexual selection is likely to be stronger compared to monogamous species (Shuster & Wade 2003). Hence, pre- and post-copulatory competition and/or mate choice seem to magnify the material that selection acts on in a sex-specific manner. In contrast, phenotypic and genetic variances in lifespan do not show sex differences in either polygamous or monogamous species. The sample size for lifespan (*N*_Estimates_ = 39; *N*_Species_ = 16) was smaller compared to reproductive success (*N*_Estimates_ = 62; *N*_Species_ = 21) meaning that we had less statistical power to detect a sex difference in phenotypic and genetic variances for lifespan. However, we did not observe any consistent trend for lifespan and our data suggest that even if there was a sex difference in genetic variance in lifespan, its effect size would be considerably smaller compared to reproductive success. Thus, we conclude that while sexual selection may promote sex differences in mean lifespan at an evolutionary scale (Lemaitre *et al*. 2020), our results indicate that it does not generate sex-specific genetic variances in this fitness component. Hence, the strength of selection on survival appears to be similar among males and females.

Despite the strong sex difference in genetic variation of reproductive success that we demonstrated, our results on the effect of the social mating system need to be considered with caution. This is because of the underrepresentation of monogamous species in our dataset and because of only two independent evolutionary changes from polygamy to monogamy in our phylogeny. Moreover, our binary classification of the mating system into socially monogamous and polygamous species fails to capture the continuum in the strength of sexual selection across the sampled taxa, likely limiting its explanatory power as a predictor variable. Ideally, one would use a standardized continuous estimate for the strength of sexual selection allowing inter- and intra-specific comparisons such as Bateman metrics (Arnold 1994; Jones 2009). Unfortunately, such data are currently only accessible for a small fraction of the sampled species (i.e. 19% based on a recent database by (Janicke & Morrow 2018)), which renders those measures as additional predictors for the strength of sexual selection unavailable at the current state of knowledge. Therefore, we argue that mating system is currently the most reliable predictor for the potential of sexual selection of the sampled species.

In essence, our findings provide support for the prediction that sexual selection promotes stronger net selection on males compared to females. We conclude that the key assumption required for sexual selection to assist natural selection and thereby to accelerate the adaptation to changing environments is often fulfilled in nature. Stronger net selection on males implies that populations may purge deleterious alleles across the genome primarily at the expense of males and thus at a low demographic cost (Agrawal 2001; Siller 2001). This has important eco-evolutionary consequences because stronger net selection on males will not only bolster local adaptation but will also reduce extinction risk when populations are coping with challenging environmental conditions (Lumley *et al*. 2015). Therefore, our findings support the idea that sexual selection can play a pivotal role in evolutionary rescue (Candolin & Heuschele 2008; Holman & Kokko 2013; Svensson 2019) and are in line with a recent meta-analysis providing compelling evidence that sexual selection increases non-sexual fitness (Cally *et al*. 2019). Interestingly, environmental stress has repeatedly been found to elevate the effect of deleterious mutations and thereby increase genetic variation in fitness-related traits (Rowinski & Rogell 2017). Thus we extend the sexes-as-environments analogy (Rice & Chippindale 2001) to say that an almost identical genome is expressed in a more stressful male environment versus a relatively more benign female environment.

Whether stronger net selection on males eventually promotes adaptation to a new environment, or even contributes to the evolution and maintenance of sexual over asexual reproduction (Agrawal 2001; Siller 2001), will also depend on another important aspect of the genetic architecture of male and female fitness components: the cross-sex genetic covariance. Specifically, only if fitness in both sexes is condition-dependent (i.e. positively affected by the amount of acquired resources) and therefore largely governed by a similar set of genes, will sexual selection on males purge deleterious alleles in females and thereby facilitate adaptation (Whitlock & Agrawal 2009). Empirical tests for an inter-specific genetic correlation of fitness revealed mixed results including examples of highly negative correlations indicating intense intra-locus sexual conflict (Foerster *et al*. 2007; Poissant *et al*. 2010). Only a small fraction of the primary studies included in our analysis reported cross-sex genetic correlations, but exploratory analyses of this subset (Supplementary Information) support an earlier finding of highly positive genetic correlations for lifespan with no consistent pattern for reproductive success (Hendry *et al*. 2018). Further work on the genetic covariance between male and female fitness components is needed to evaluate the overall potential of sexual selection to facilitate or constrain the adaptation to changing environments. Specifically, how genetic variances and covariances of male and female fitness components change with ecological conditions is largely unknown (Delcourt *et al*. 2009; Berger *et al*. 2014) though such knowledge is crucial to predict evolutionary trajectories when populations face environmental change. Moreover, for some taxa, sexual selection has been found to increase extinction risk (Doherty *et al*. 2003; Le Galliard *et al*. 2005) potentially as a consequence of intense sexual conflict but the quantitative genetics of male and female fitness of those species remain mostly unexplored.

Despite the detected sex differences and the effect of the social mating system, a large fraction of the intra- and inter-specific variance in *CV*_G_ remained unexplained (Table 1). This is potentially, at least in part, because genetic variances are often estimated with low precision, which may have introduced substantial noise into our analyses. Besides that, we suspect that another part of the unexplained variation stems from environmental effects influencing genetic (co)variances (i.e., genotype by environmental interactions) (Rowinski & Rogell 2017), which limits comparisons of studies conducted under different experimental conditions. Moreover, there are also methodological differences between primary studies, which may have contributed to the unexplained variation in *CV*_G_. This includes the application of different breeding designs used to quantify genetic variances (i.e. pedigrees, half-sib/full-sib breeding, parent-offspring regressions, inbred lines), differences in the analytical approaches (latent-scale versus data-scale estimates of genetic variances), and disparity in the measurement of reproductive success (annual versus lifetime reproductive success; see Material and Methods). While all these sources of uncontrolled variation are likely to have introduced noise into our dataset, we are not aware of any systematic biases that they might have created. Finally, our study covers a broad taxonomic range spanning flatworms, mollusks, arthropods, and vertebrates but is based on relatively few species. This is admittedly a limitation of our study but at the same time illustrates the clear need for more quantitative genetic studies measuring genetic variation of fitness components in both sexes.

Collectively, our analysis highlights the role that sexual selection has for generating sex differences in the strength of net selection. However, we have just started to understand the eco-evolutionary consequences of sexual selection in terms of its impact on demography and the adaptive potential of populations to cope with changing environments.

## Acknowledgements

We are very grateful to all authors of the primary studies and in particular those providing additional information and/or data including Jessica Abbott, Mats Björklund, Sandra Bouwhuis, Ryan G. Calsbeek, Julie Collet, David Hosken, Zenobia Lewis, Jacob Moorad, Tom Tregenza and Felix Zajitschek. Moreover, we thank Patrice David, Shinichi Nakagawa, Klaus Reinhardt, Holger Schielzeth, Céline Teplitsky and Pierre de Villemereuil for discussions and statistical advice.

## Supplementary Information

**Fig. S1.**
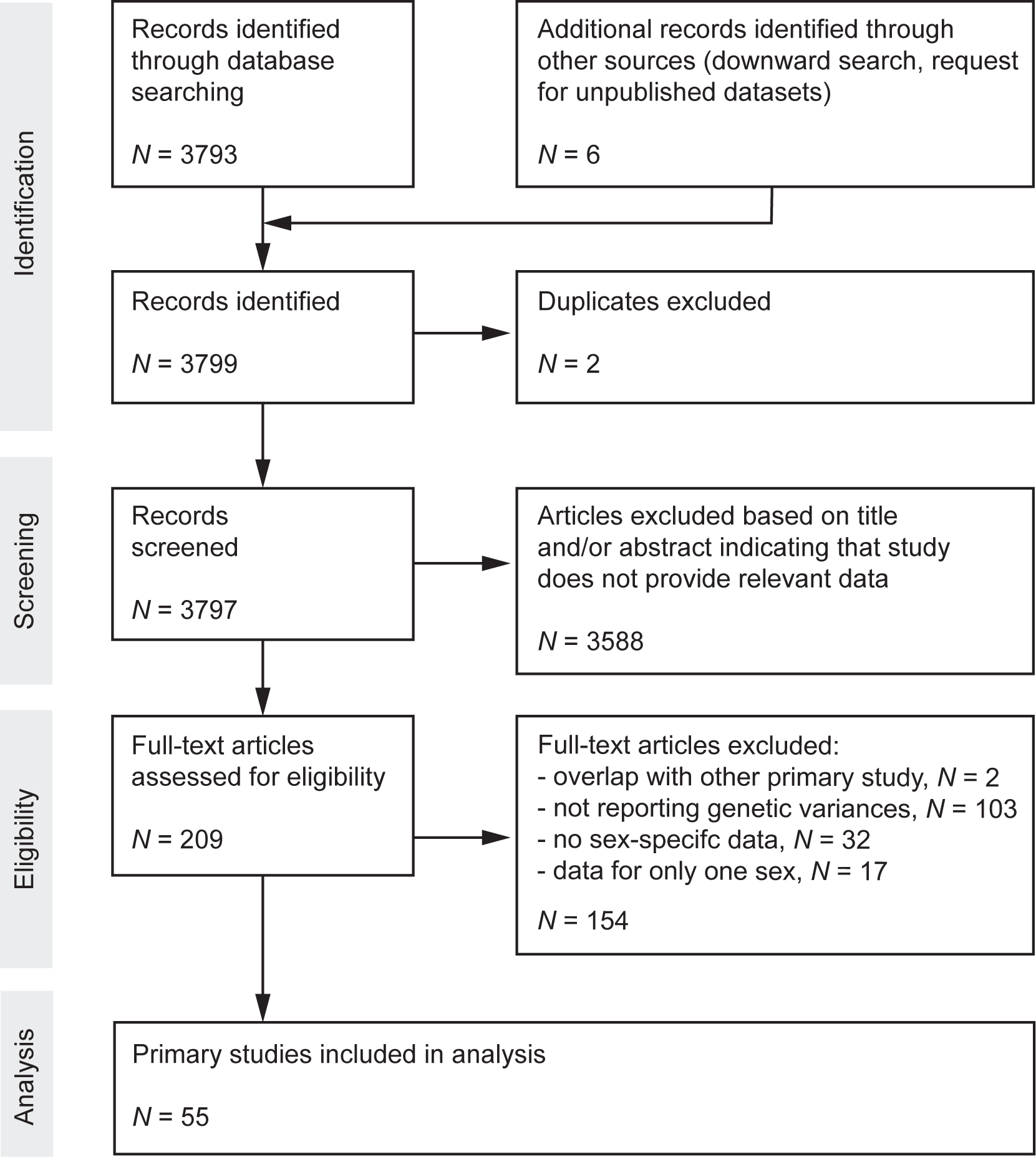
Preferred Reporting Items for Systematic Reviews and Meta-Analyses (PRISMA) Diagram. Flow chart maps the number of records identified during the different phases of the systematic literature search.

**Fig. S2.**
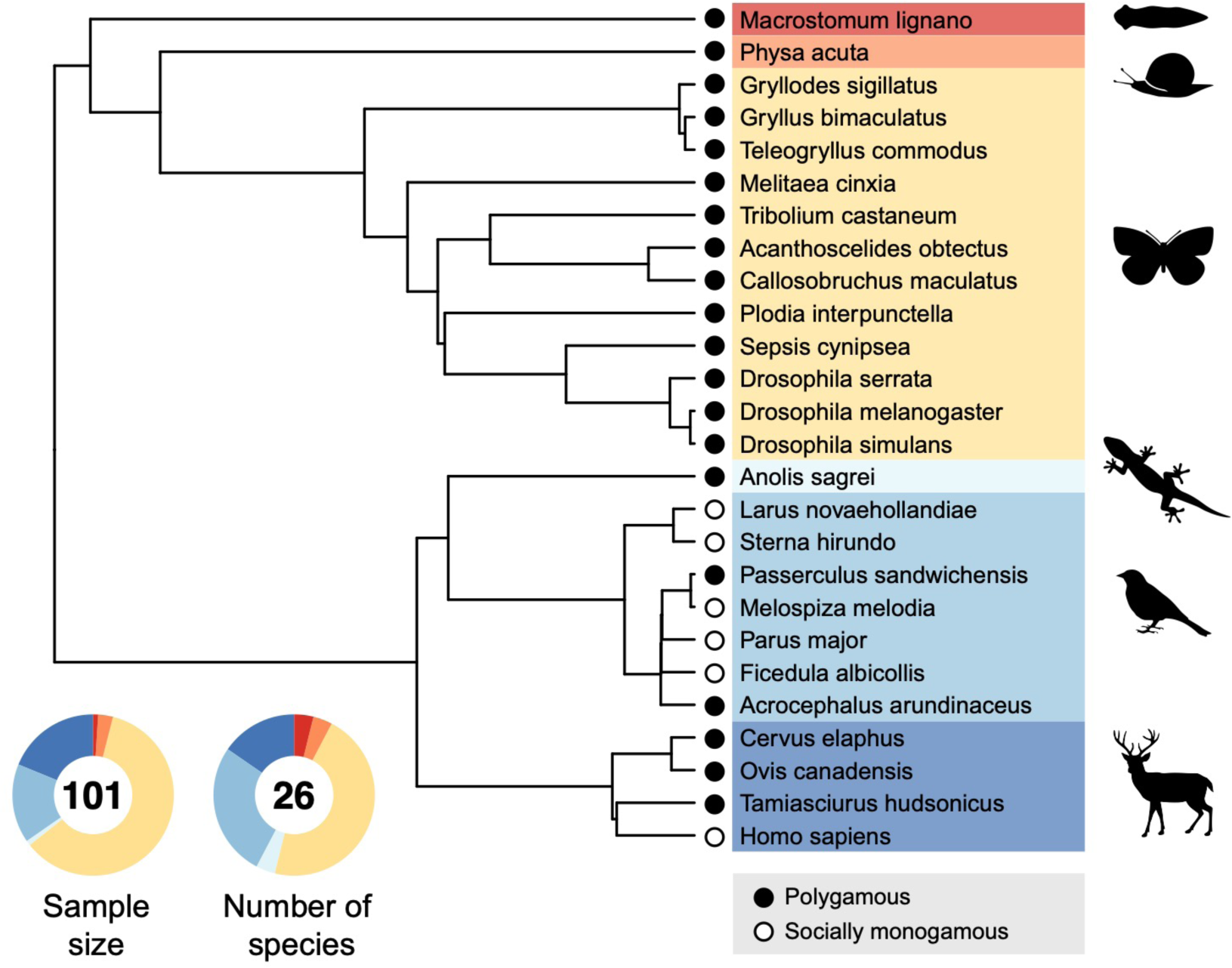
Phylogeny used to account for phylogenetic non-independence in statistical modelling. Doughnut charts show the relative fraction of the sample size (i.e. number of paired estimates for male and female genetic variance) and the number of species, both pooled for reproductive success and lifespan.

**Fig. S3.**
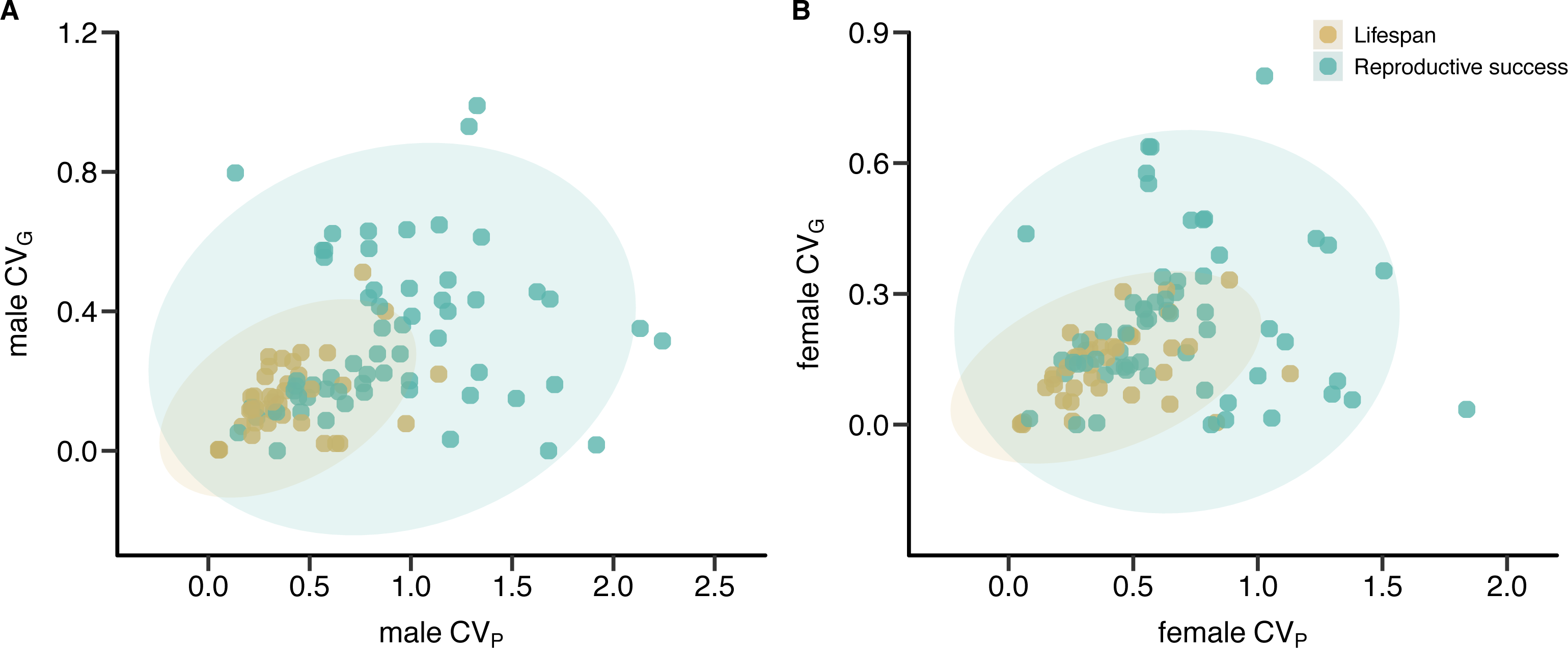
Positive correlation between phenotypic and genetic coefficients of variation (*CV*_P_ and *CV*_G_, respectively) for lifespan but not for reproductive success. Scatterplots show relationships between *CV*_P_ and *CV*_G_ for male (A) and female (B) reproductive success (green) and lifespan (brown). Shaded areas indicate the 95 % confidence ellipses.

**Table S1.** Results of PGLMMs testing for sex by mating system interaction on phenotypic (*CV_P_*) and genetic (*CV_G_*) coefficient of variation. Results shown for reproductive success (RS) and lifespan (LS). Estimates are shown as posterior means with 95% Highest Posterior Density (HPD) intervals.

| Response | Variance component | Predictor | Estimate | $P_{MCMC}$ |
| --- | --- | --- | --- | --- |
| RS | $CV_P$ | Sex | -0.012 (-0.172, 0.149) | 0.886 |
|  |  | Mating system | -0.001 (-0.378, 0.386) | 0.968 |
|  |  | Sex by Mating system | 0.324 (0.138, 0.508) | < 0.001 |
| | $CV_G$ | Sex | -0.015 (-0.101, 0.065) | 0.730 |
|  |  | Mating system | -0.005 (-0.155, 0.136) | 0.948 |
|  |  | Sex by Mating system | 0.134 (0.040, 0.231) | 0.007 |
| LS | $CV_P$ | Sex | 0.026 (-0.025, 0.081) | 0.325 |
|  |  | Mating system | 0.198 (-0.115, 0.512) | 0.208 |
|  |  | Sex by Mating system | -0.032 (-0.090, 0.032) | 0.296 |
| | $CV_G$ | Sex | 0.050 (0.004, 0.094) | 0.029 |
|  |  | Mating system | -0.028 (-0.147, 0.089) | 0.618 |
|  |  | Sex by Mating system | -0.041 (-0.091, 0.011) | 0.124 |

**Table S2.** Phylogenetic independent meta-analysis of lnCVR defined as the log-transformed ratio of male to female coefficient of phenotypic variation. Models are shown testing for (a) the intercept across mating systems (i.e. global effect size with positive values indicating a male bias), (b) with mating system as moderator variable and separately for (c) monogamous and (d) polygamous species. Results shown for reproductive success (RS) and lifespan (LS). Estimates are shown as posterior means with 95% Highest Posterior Density (HPD) intervals.

| Response | Predictor | Estimate | | $P_{\text{MCMC}}$ |
| --- | --- | --- | --- | --- |
| (a) Across mating systems |  |  |  |  |
| RS | Intercept | 0.232 | (-0.249, 0.705) | 0.257 |
| LS | Intercept | 0.030 | (-0.094, 0.151) | 0.576 |
| (b) Mating system |  |  |  |  |
| RS | Intercept | -0.032 | (-0.360, 0.266) | 0.835 |
|  | Mating system | 0.363 | (0.124, 0.624) | 0.011 |
| LS | Intercept | 0.082 | (-0.107, 0.275) | 0.357 |
|  | Mating system | -0.077 | (-0.270, 0.129) | 0.436 |
| (c) Monogamy |  |  |  |  |
| RS | Intercept | 0.056 | (-1.336, 1.470) | 0.791 |
| LS | Intercept | 0.064 | (-0.325, 0.454) | 0.508 |
| (d) Polygamy |  |  |  |  |
| RS | Intercept | 0.326 | (0.007, 0.612) | 0.043 |
| LS | Intercept | 0.006 | (-0.166, 0.192) | 0.927 |

**Table S3.** Results of PGLMMs testing for the effect of sex, study type (lab *versus* field studies) and their interaction on phenotypic (*CV_P_*) and genetic (*CV_G_*) coefficient of variation. Results shown for reproductive success (RS) and lifespan (LS). Estimates are shown as posterior means with 95% Highest Posterior Density (HPD) intervals.

| Response | Variance component | Predictor | Estimate | $P_{MCMC}$ |
| --- | --- | --- | --- | --- |
| RS | $CV_P$ | Sex | 0.187 (0.044, 0.328) | 0.012 |
|  |  | Study type | -0.308 (-0.661, 0.028) | 0.074 |
|  |  | Sex by Study type | 0.074 (-0.103, 0.253) | 0.418 |
| | $CV_G$ | Sex | 0.084 (0.010, 0.153) | 0.020 |
|  |  | Study type | 0.124 (-0.154, 0.401) | 0.302 |
|  |  | Sex by Study type | 0.004 (-0.085, 0.094) | 0.934 |
| LS | $CV_P$ | Sex | -0.003 (-0.046, 0.042) | 0.883 |
|  |  | Study type | -0.229 (-0.644, 0.206) | 0.183 |
|  |  | Sex by Study type | -0.002 (-0.057, 0.051) | 0.938 |
| | $CV_G$ | Sex | 0.024 (-0.013, 0.063) | 0.204 |
|  |  | Study type | 0.023 (-0.252, 0.334) | 0.856 |
|  |  | Sex by Study type | -0.012 (-0.057, 0.036) | 0.618 |

### Supplementary Analysis

#### Cross-sex genetic correlations

Theory predicting a positive effect of sexual selection on adaptation is based on the additional assumption that alleles favored in males must also be beneficial in females, which, if true, may manifest in a positive genetic correlation between male and female fitness. Estimates of such cross-sex genetic correlations are scarce and a previous comparative study included only few correlation of fitness components relative to morphological, physiological and behavioral traits (Poissant *et al*. 2010).

In an explorative meta-analysis, we tested for an overall positive genetic cross-sex correlation in reproductive success and lifespan based on estimates provided in the primary studies obtained from our systematic literature search (Fig. S1). Only few primary studies reported cross-sex correlations, which led to small samples sizes for reproductive success (N = 25) and lifespan (N = 19), and is also reflected in low numbers of sampled species (reproductive success: *N*_Species_ = 10; lifespan: *N*_Species_ = 5).

We used phylogenetically informed meta-analyses by running PGLMMs with the *MCMCglmm* R package (using same specifications as for models reported in the main text), in which Pearson correlation coefficients (*r*) were defined as response variable weighted by the inverse of its variance. Models included study identifier and phylogeny as random effect terms. We found support for positive cross-sex genetic correlation for lifespan (PGLMM: global *r* = 0.548, 95% CI = 0.185 – 0.918, *P* = 0.020) but not for reproductive success (PGLMM: global *r* = 0.024, 95% CI = −0.242 – 0.244, *P* = 0.816). In conclusion, our limited data does not support a positive cross-sex genetic correlation for reproductive success.

### Supplementary Data

#### Mating system classification

**Table S5.** Mating system classification (monogamy versus polygamy) of the 26 sampled species in alphabetical order.

| Species | Class | Mating System | Reference |
| --- | --- | --- | --- |
| *Acanthoscelides obtectus* | Insecta | Polygamy | (Seslija *et al.* 2009) |
| *Acrocephalus arundinaceus* | Aves | Polygamy | (Hasselquist *et al.* 1995) |
| *Anolis sagrei* | Reptilia | Polygamy | (Kamath & Losos 2018) |
| *Callosobruchus maculatus* | Insecta | Polygamy | (Fritzsche & Arnqvist 2013) |
| *Cervus elaphus* | Mammalia | Polygamy | (Bonenfant *et al.* 2004; Frantz *et al.* 2008) |
| *Drosophila melanogaster* | Insecta | Polygamy | (Bjork & Pitnick 2006; Morimoto *et al.* 2016) |
| *Drosophila serrata* | Insecta | Polygamy | (Frentiu & Chenoweth 2008) |
| *Drosophila simulans* | Insecta | Polygamy | (Taylor *et al.* 2008) |
| *Ficedula albicollis* | Aves | Monogamy | (Qvarnstrom *et al.* 2003) |
| *Gryllodes sigillatus* | Insecta | Polygamy | (Ivy & Sakaluk 2005) |
| *Gryllus bimaculatus* | Insecta | Polygamy | (Bretman & Tregenza 2005) |
| *Homo sapiens* | Mammalia | Monogamy | (Dixson 2009) |
| *Larus novaehollandiae* | Aves | Monogamy | (Mills 1973) |
| *Macrostomum lignano* | Rhabditophora | Polygamy | (Janicke & Schärer 2009) |
| *Melitaea cinxia* | Insecta | Polygamy | (Boggs & Nieminen 2004) |
| *Melospiza melodia* | Aves | Monogamy | (Arcese 1989) |
| *Ovis canadensis* | Mammalia | Polygamy | (Coltman *et al.* 2002) |
| *Parus major* | Aves | Monogamy | (Dhondt 1987) |
| *Passerculus sandwichensis* | Aves | Polygamy | (Wheelwright *et al.* 1992) |
| *Physa acuta* | Gastropoda | Polygamy | (Pélissié *et al.* 2012) |
| *Plodia interpunctella* | Insecta | Polygamy | (Gage 1995) |
| *Sepsis cynipsea* | Insecta | Polygamy | (Ward 1983; Teuschl & Blanckenhorn 2007) |
| *Sterna hirundo* | Aves | Monogamy | (Griggio *et al.* 2004) |
| *Tamiasciurus hudsonicus* | Mammalia | Polygamy | (Lane *et al.* 2008) |
| *Teleogryllus commodus* | Insecta | Polygamy | (Evans 1988; Jennions *et al.* 2004) |
| *Tribolium castaneum* | Insecta | Polygamy | (Fedina & Lewis 2008) |

### Supplementary Data

#### Search terms and list of primary studies

Systematic literature search was carried out in the Web of Science Core Collection (Clarivate) using the following search terms:

TS=((sex* OR (male AND female) OR (man AND woman) OR “sex diff*” OR “gender diff*” OR sex-specific OR intersex* OR inter-sex* OR cross-sex* OR “across sex*” OR “between sex*” OR “between-sex*” OR “sex-limited”) AND (fitness OR “reproductive success” OR survival OR longevity OR lifespan OR “life span”) AND (“intra-locus sexual conflict” OR “intralocus sexual conflict” OR “sexually antagonistic genetic” OR “genetic co*” OR heritability OR “genetic varia*” OR “quantitative genetics” OR “genetic architecture” OR evolvability))

The list below encompasses all 52 published primary studies included in the comparative analyses. It does not comprise three unpublished studies that have also been included (Abbott, J., and A. Norden. in prep.; Janicke, T., E. Chapuis, S. Meconcelli, N. Bonel, and P. David. in prep.; Moiron M, Charmantier A, Bouwhuis S. in prep.).

## References

Agrawal, A.F. (2001). Sexual selection and the maintenance of sexual reproduction. Nature, 411, 692–695.

Andersson, M. (1994). Sexual Selection. Princeton University Press, Princeton.

Arnold, S.J. (1994). Bateman principles and the measurement of sexual selection in plants and animals. Am. Nat., 144, S126–S149.

Arnqvist, G. & Rowe, L. (2005). Sexual Conflict. Princeton University Press, Princeton, NJ, USA.

Bateman, A.J. (1948). Intra-sexual selection in *Drosophila*. Heredity, 2, 349–368.

Berger, D., Grieshop, K., Lind, M.I., Goenaga, J., Maklakov, A.A. & Arnqvist, G. (2014). Intralocus sexual conflict and environmental stress. Evolution, 68, 2184–2196.

Bolund, E., Bouwhuis, S., Pettay, J.E. & Lummaa, V. (2013). Divergent selection on, but no genetic conflict over, female and male timing and rate of reproduction in a human population. Proc. R. Soc. B-Biol. Sci., 280, 9.

Bonduriansky, R., Maklakov, A., Zajitschek, F. & Brooks, R. (2008). Sexual selection, sexual conflict and the evolution of ageing and life span. Funct. Ecol., 22, 443–453.

Brouwer, L. & Griffith, S.C. (2019). Extra-pair paternity in birds. Mol. Ecol., 28, 4864–4882.

Cally, J.G., Stuart-Fox, D. & Holman, L. (2019). Meta-analytic evidence that sexual selection improves population fitness. Nature Communications, 10, 10.

Candolin, U. & Heuschele, J. (2008). Is sexual selection beneficial during adaptation to environmental change? Trends Ecol. Evol., 23, 446–452.

Chippindale, A.K., Gibson, J.R. & Rice, W.R. (2001). Negative genetic correlation for adult fitness between sexes reveals ontogenetic conflict in *Drosophila*. Proc. Natl. Acad. Sci. U. S. A., 98, 1671–1675.

Clutton-Brock, T. (2007). Sexual selection in males and females. Science, 318, 1882–1885.

Connallon, T. & Matthews, G. (2019). Cross-sex genetic correlations for fitness and fitness components: Connecting theoretical predictions to empirical patterns. Evol. Lett., 3, 254–262.

Cotton, S., Fowler, K. & Pomiankowski, A. (2004). Do sexual ornaments demonstrate heightened condition-dependent expression as predicted by the handicap hypothesis? Proceedings of the Royal Society B: Biological Sciences, 271, 771– 783.

Crow, J.F. (1958). Some possibilities for measuring selection intensities in man. Human Biology, 30, 1–13.

de Villemereuil, P. & Nakagawa, S. (2014). General quantitative genetic methods for comparative biology. In: Modern phylogenetic comparative methods and their application in evolutionary biology (ed. Garamszegi, LZ). Springer-Verlag Berlin Heidelberg, p. 552.

Delcourt, M., Blows, M.W. & Rundle, H.D. (2009). Sexually antagonistic genetic variance for fitness in an ancestral and a novel environment. Proc. R. Soc. B-Biol. Sci., 276, 2009–2014.

Doherty, P.F., Sorci, G., Royle, J.A., Hines, J.E., Nichols, J.D. & Boulinier, T. (2003). Sexual selection affects local extinction and turnover in bird communities. Proc. Natl. Acad. Sci. U. S. A., 100, 5858–5862.

Dunn, P.O., Whittingham, L.A. & Pitcher, T.E. (2001). Mating systems, sperm competition, and the evolution of sexual dimorphism in birds. Evolution, 55, 161–175.

Fisher, R.A. (1930). The Genetical Theory of Natural Selection. Oxford University Press., Oxford, UK.

Foerster, K., Coulson, T., Sheldon, B.C., Pemberton, J.M., Clutton-Brock, T.H. & Kruuk, L.E. (2007). Sexually antagonistic genetic variation for fitness in red deer. Nature, 447, 1107–1110.

Fox, C., Bush, M., Roff, D. & Wallin, W. (2004). Evolutionary genetics of lifespan and mortality rates in two populations of the seed beetle, Callosobruchus maculatus. Heredity, 92, 170–181.

Gay, L., Brown, E., Tregenza, T., Pincheira-Donoso, D., Eady, P.E., Vasudev, R. et al. (2011). The genetic architecture of sexual conflict: male harm and female resistance in Callosobruchus maculatus. J. Evol. Biol., 24, 449–456.

Grafen, A. (1991). Modelling in behavioural ecology. In: Behavioural Ecology (eds. Krebs, JR & Davies, NB). Blackwell Scientific Publications Oxford, pp. 5–31.

Hadfield, J.D. (2010). MCMC methods for multi-response generalized linear mixed models: the MCMCglmm R package. Journal of Statistical Software, 33, 1–22.

Haldane, J.B.S. (1937). The effect of variation on fitness. Am. Nat., 71, 337–349.

Hendry, A.P., Schoen, D.J., Wolak, M.E. & Reid, J.M. (2018). The contemporary evolution of fitness. Annual Review of Ecology, Evolution, and Systematics, 49, 457–476.

Holman, L. & Jacomb, F. (2017). The effects of stress and sex on selection, genetic covariance, and the evolutionary response. J. Evol. Biol., 30, 1898–1909.

Holman, L. & Kokko, H. (2013). The consequences of polyandry for population viability, extinction risk and conservation. Philos. Trans. R. Soc. B-Biol. Sci., 368, 12.

Houle, D. (1992). Comparing evolvability and variation of quantitative traits. Genetics, 130, 195–204.

Janicke, T., Häderer, I.K., Lajeunesse, M.J. & Anthes, N. (2016). Darwinian sex roles confirmed across the animal kingdom. Science Advances, 2, e1500983.

Janicke, T. & Morrow, E.H. (2018). Operational sex ratio predicts the opportunity and direction of sexual selection across animals. Ecol. Lett., 21, 384–391

Jennions, M.D., Moller, A.P. & Petrie, M. (2001). Sexually selected traits and adult survival: A meta-analysis. Q. Rev. Biol., 76, 3–36.

Jones, A.G. (2009). On the opportunity for sexual selection, the Bateman gradient and the maximum intensity of sexual selection. Evolution, 63, 1673–1684.

Kumar, S., Stecher, G., Li, M., Knyaz, C. & Tamura, K. (2018). MEGA X: molecular evolutionary genetics analysis across computing platforms. Mol. Biol. Evol., 35, 1547–1549.

Kumar, S., Stecher, G., Suleski, M. & Hedges, S.B. (2017). TimeTree: A Resource for Timelines, Timetrees, and Divergence Times. Mol. Biol. Evol., 34, 1812–1819.

Le Galliard, J.F., Fitze, P.S., Ferriere, R. & Clobert, J. (2005). Sex ratio bias, male aggression, and population collapse in lizards. Proc. Natl. Acad. Sci. U. S. A., 102, 18231–18236.

Lemaitre, J.F., Ronget, V., Tidiere, M., Allaine, D., Berger, V., Cohas, A. et al. (2020). Sex differences in adult lifespan and aging rates of mortality across wild mammals. Proc. Natl. Acad. Sci. U. S. A., 117, 8546–8553.

Lorch, P.D., Proulx, S., Rowe, L. & Day, T. (2003). Condition-dependent sexual selection can accelerate adaptation. Evol. Ecol. Res., 5, 867–881.

Lukas, D. & Clutton-Brock, T.H. (2013). The Evolution of Social Monogamy in Mammals. Science, 341, 526–530.

Lumley, A.J., Michalczyk, L., Kitson, J.J.N., Spurgin, L.G., Morrison, C.A., Godwin, J.L. et al. (2015). Sexual selection protects against extinction. Nature, 522, 470–473.

Macartney, E.L., Crean, A.J., Nakagawa, S. & Bonduriansky, R. (2019). Effects of nutrient limitation on sperm and seminal fluid: a systematic review and meta-analysis. Biol. Rev., 94, 1722–1739.

Martínez-Ruiz, C. & Knell, R.J. (2017). Sexual selection can both increase and decrease extinction probability: reconciling demographic and evolutionary factors. J. Anim. Ecol., 86, 117–127.

Martinossi-Allibert, I., Rueffler, C., Arnqvist, G. & Berger, D. (2019). The efficacy of good genes sexual selection under environmental change. Proc. R. Soc. B-Biol. Sci., 286, 9.

Moller, A.P. (1986). Mating systems among European passerines: a review. Ibis, 128, 234–250.

Nakagawa, S., Poulin, R., Mengersen, K., Reinhold, K., Engqvist, L., Lagisz, M. et al. (2015). Meta-analysis of variation: ecological and evolutionary applications and beyond. Methods in Ecology and Evolution, 6, 143–152.

Nakagawa, S. & Schielzeth, H. (2013). A general and simple method for obtaining R2 from generalized linear mixed-effects models. Methods in Ecology and Evolution, 4, 133–142.

Paradis, E. & Schliep, K. (2019). ape 5.0: an environment for modern phylogenetics and evolutionary analyses in R. Bioinformatics, 35, 526–528.

Pélissié, B., Jarne, P. & David, P. (2012). Sexual selection without sexual dimorphism: Bateman gradients in a simultaneous hermaphrodite. Evolution, 66, 66–81.

Poissant, J., Wilson, A.J. & Coltman, D.W. (2010). Sex-specific genetic variance and the evolution of sexual Dimorphism: a systematic review of cross-sex genetic correlations. Evolution, 64, 97–107.

Rice, W.R. & Chippindale, A.K. (2001). Intersexual ontogenetic conflict. J. Evol. Biol., 14, 685–693.

Rowe, L. & Houle, D. (1996). The lek paradox and the capture of genetic variance by condition dependent traits. Proc. R. Soc. B-Biol. Sci., 263, 1415–1421.

Rowinski, P.K. & Rogell, B. (2017). Environmental stress correlates with increases in both genetic and residual variances: A meta-analysis of animal studies. Evolution, 71, 1339–1351.

Shuster, S.M. & Wade, M.J. (2003). Mating Systems and Strategies. Princeton University Press.

Siller, S. (2001). Sexual selection and the maintenance of sex. Nature, 411, 689–692.

Svensson, E.I. (2019). Eco-evolutionary dynamics of sexual selection and sexual conflict. Funct. Ecol., 33, 60–72.

Webb, C.O., Ackerly, D.D. & Kembel, S.W. (2008). Phylocom: software for the analysis of phylogenetic community structure and trait evolution. Bioinformatics, 24, 2098–2100.

Wheelwright, N.T., Keller, L.F. & Postma, E. (2014). The effect of trait type and strength of selection on heritability and evolvability in an island bird population. Evolution, 68, 3325–3336.

Whitlock, M.C. & Agrawal, A.F. (2009). Purging the genome with sexual selection: reducing mutation load through selection on males. Evolution, 63, 569–582.

## References – Mating system classification

Arcese, P. (1989). Intrasexual competition, mating system and natal dispersal in song sparrows. Anim. Behav., 38, 958–979.

Bjork, A. & Pitnick, S. (2006). Intensity of sexual selection along the anisogamy-isogamy continuum. Nature, 441, 742–745.

Boggs, C. & Nieminen, M.J. (2004). Cherckerspot reproductive biology. In: On the wings of checkerspots: A model system for population biology. Oxford University Press, pp. 92–111.

Bonenfant, C., Gaillard, J.M., Klein, F. & Maillard, D. (2004). Variation in harem size of red deer (*Cervus elaphus* L.): the effects of adult sex ratio and age-structure. J. Zool., 264, 77–85.

Bretman, A. & Tregenza, T. (2005). Measuring polyandry in wild populations: a case study using promiscuous crickets. Mol. Ecol., 14, 2169–2179.

Coltman, D.W., Festa-Bianchet, M., Jorgenson, J.T. & Strobeck, C. (2002). Age-dependent sexual selection in bighorn rams. Proc. R. Soc. B-Biol. Sci., 269, 165–172.

Dhondt, A.A. (1987). Polygynous blue tits and monogamous great tits: does the polygyny-threshold model hold? Am. Nat., 129, 213–220.

Dixson, A.F. (2009). Sexual selection and the origins of human mating systems. Oxford University Press.

Evans, A. (1988). Mating systems and reproductive strategies in three australian gryllid crickets: *Bobilla victoriae* Otte, *Balamara gidya* Otte and *Teleogryllus commodus* (Walker) (Orthoptera: Gryllidae: Nemobiinae; Trigonidiinae; Gryllinae). Ethology.

Fedina, T.Y. & Lewis, S.M. (2008). An integrative view of sexual selection in Tribolium flour beetles. Biol. Rev., 83, 151–171.

Frantz, A.C., Hamann, J.L. & Klein, F. (2008). Fine-scale genetic structure of red deer (*Cervus elaphus*) in a French temperate forest. European Journal of Wildlife Research, 54, 44–52.

Frentiu, F.D. & Chenoweth, S.F. (2008). Polyandry and paternity skew in natural and experimental populations of *Drosophila serrata*. Mol. Ecol., 17, 1589–1596.

Fritzsche, K. & Arnqvist, G. (2013). Homage to Bateman: sex roles predict sex differences in sexual selection. Evolution; international journal of organic evolution, 67, 1926–1936.

Gage, M.J. (1995). Continuous variation in reproductive strategy as an adaptive response to population density in the moth *Plodia interpunctella*. Proceedings of the Royal Society of London. Series B: Biological Sciences, 261, 25–30.

Griggio, M., Matessi, G. & Marin, G. (2004). No evidence of extra-pair paternity in a colonial seabird, the common tern (*Sterna hirundo*). *Ital*. J. Zool., 71, 219–222.

Hasselquist, D., Bensch, S. & Vonschantz, T. (1995). Low frequency of extrapair paternity in the polygynous great reed warbler, *Acrocephalus arundinaceus*. Behav. Ecol., 6, 27–38.

Ivy, T.M. & Sakaluk, S.K. (2005). Polyandry promotes enhanced offspring survival in decorated crickets. Evolution, 59, 152–159.

Janicke, T. & Schärer, L. (2009). Determinants of mating and sperm-transfer success in a simultaneous hermaphrodite. Journal of Evolutionary Biology, 22, 405–415.

Jennions, M.D., Hunt, J., Graham, R. & Brooks, R. (2004). No evidence for inbreeding avoidance through postcopulatory mechanisms in the black field cricket, *Teleogryllus commodus*. Evolution, 58, 2472–2477.

Kamath, A. & Losos, J.B. (2018). Estimating encounter rates as the first step of sexual selection in the lizard *Anolis sagrei*. Proc. R. Soc. B-Biol. Sci., 285, 9.

Lane, J.E., Boutin, S., Gunn, M.R., Slate, J. & Coltman, D.W. (2008). Female multiple mating and paternity in free-ranging North American red squirrels. Anim. Behav., 75, 1927–1937.

Mills, J.A. (1973). The influence of age and pair-bond on the breeding biology of the red-billed gull *Larus novaehollandiae scopulinus*. J. Anim. Ecol., 42, 147–162.

Morimoto, J., Pizzari, T. & Wigby, S. (2016). Developmental environment effects on sexual selection in male and female *Drosophila melanogaster*. PLoS One, 11, 27.

Qvarnstrom, A., Sheldon, B.C., Part, T. & Gustafsson, L. (2003). Male ornamentation, timing of breeding, and cost of polygyny in the collared flycatcher. Behav. Ecol., 14, 68–73.

Seslija, D., Lazarevic, J., Jankovic, B. & Tucic, N. (2009). Mating behavior in the seed beetle *Acanthoscelides obtectus* selected for early and late reproduction. Behav. Ecol., 20, 547–552.

Taylor, M.L., Wigmore, C., Hodgson, D.J., Wedell, N. & Hosken, D.J. (2008). Multiple mating increases female fitness in *Drosophila simulans*. Anim. Behav., 76, 963–970.

Teuschl, Y. & Blanckenhorn, W.U. (2007). The reluctant fly: what makes *Sepsis cynipsea* females willing to copulate? Anim. Behav., 73, 85–97.

Ward, P.I. (1983). The effects of size on the mating behaviour of the dung fly *Sepsis cynipsea*. Behav. Ecol. Sociobiol., 13, 75–80.

Wheelwright, N.T., Schultz, C.B. & Hodum, P.J. (1992). Polygyny and male parental care in Savannah sparrows: effects on female fitness. Behav. Ecol. Sociobiol., 31, 279–289.

## References

Archer, C. R., F. Zajitschek, S. K. Sakaluk, N. J. Royle, and J. Hunt. 2012. Sexual selection affects the evolution of lifespan and ageing in the decorated cricket *Gryllodes sigillatus*. Evolution 66:3088–3100.

Berger, D., K. Grieshop, M. I. Lind, J. Goenaga, A. A. Maklakov, and G. Arnqvist. 2014. Intralocus sexual conflict and environmental stress. Evolution 68:2184–2196.

Bolund, E., S. Bouwhuis, J. E. Pettay, and V. Lummaa. 2013. Divergent selection on, but no genetic conflict over, female and male timing and rate of reproduction in a human population. Proceedings of the Royal Society B-Biological Sciences 280.

Brommer, J. E., M. Kirkpatrick, A. Qvarnstrom, and L. Gustafsson. 2007. The intersexual genetic correlation for lifetime fitness in the wild and its implications for sexual selection. Plos One 2.

Calsbeek, R., M. C. Duryea, D. Goedert, P. Bergeron, and R. M. Cox. 2015. Intralocus sexual conflict, adaptive sex allocation, and the heritability of fitness. Journal of Evolutionary Biology 28:1975–1985.

Collet, J. M., S. Fuentes, J. Hesketh, M. S. Hill, P. Innocenti, E. H. Morrow, K. Fowler et al. 2016. Rapid evolution of the intersexual genetic correlation for fitness in *Drosophila melanogaster*. Evolution 70:781–795.

Coltman, D. W., P. O’Donoghue, J. T. Hogg, and M. Festa-Bianchet. 2005. Selection and genetic (co)variance in bighorn sheep. Evolution 59:1372–1382.

Delcourt, M., M. W. Blows, and H. D. Rundle. 2009. Sexually antagonistic genetic variance for fitness in an ancestral and a novel environment. Proceedings of the Royal Society B-Biological Sciences 276:2009–2014.

Duffy, E., C. R. Archer, M. D. Sharma, M. Prus, R. A. Joag, J. Radwan, N. Wedell et al. 2019. Wolbachia infection can bias estimates of intralocus sexual conflict. Ecology and Evolution 9:328–338.

Duffy, E., R. Joag, J. Radwan, N. Wedell, and D. J. Hosken. 2014. Inbreeding alters intersexual fitness correlations in *Drosophila simulans*. Ecology and Evolution 4:3330–3338.

Foerster, K., T. Coulson, B. C. Sheldon, J. M. Pemberton, T. H. Clutton-Brock, and L. E. B. Kruuk. 2007. Sexually antagonistic genetic variation for fitness in red deer. Nature 447:1107–U1109.

Fox, C. W., M. L. Bush, D. A. Roff, and W. G. Wallin. 2004. Evolutionary genetics of lifespan and mortality rates in two populations of the seed beetle, *Callosobruchus maculatus*. Heredity 92:170–181.

Gavrus-Ion, A., T. Sjovold, M. Hernandez, R. Gonzalez-Jose, M. E. E. Torne, N. Martinez-Abadias, and M. Esparza. 2017. Measuring fitness heritability: Life history traits versus morphological traits in humans. American Journal of Physical Anthropology 164:321–330.

Gay, L., E. Brown, T. Tregenza, D. Pincheira-Donoso, P. E. Eady, R. Vasudev, J. Hunt et al. 2011. The genetic architecture of sexual conflict: male harm and female resistance in *Callosobruchus maculatus*. Journal of Evolutionary Biology 24:449–456.

Griffin, R. M., H. Schielzeth, and U. Friberg. 2016. Autosomal and X-linked additive genetic variation for lifespan and aging: Comparisons within and between the sexes in *Drosophila melanogaster*. G3-Genes Genomes Genetics 6:3903–3911.

Hallsson, L. R., and M. Bjorklund. 2012. Sex-specific genetic variances in life-history and morphological traits of the seed beetle *Callosobruchus maculatus*. Ecology and Evolution 2:128–138.

Holman, L., and F. Jacomb. 2017. The effects of stress and sex on selection, genetic covariance, and the evolutionary response. Journal of Evolutionary Biology 30:1898–1909.

Innocenti, P., and E. H. Morrow. 2010. The sexually antagonistic genes of *Drosophila melanogaster*. Plos Biology 8.

Kimber, C. M., and A. K. Chippindale. 2013. Mutation, condition, and the maintenance of extended lifespan in *Drosophila*. Current Biology 23:2283–2287.

Klemme, I., and I. Hanski. 2009. Heritability of and strong single gene (Pgi) effects on life-history traits in the Glanville fritillary butterfly. Journal of Evolutionary Biology 22:1944–1953.

Kohler, H. P., J. L. Rodgers, and K. Christensen. 1999. Is fertility behavior in our genes? Findings from a Danish twin study. Population and Development Review 25:253-+.

Kosova, G., M. Abney, and C. Ober. 2010. Heritability of reproductive fitness traits in a human population. Proceedings of the National Academy of Sciences of the United States of America 107:1772–1778.

Kruuk, L. E. B., T. H. Clutton-Brock, J. Slate, J. M. Pemberton, S. Brotherstone, and F. E. Guinness. 2000. Heritability of fitness in a wild mammal population. Proceedings of the National Academy of Sciences of the United States of America 97:698–703.

Lehtovaara, A., H. Schielzeth, I. Flis, and U. Friberg. 2013. Heritability of life span is largely sex limited in *Drosophila*. American Naturalist 182:653–665.

Leips, J., and T. F. C. Mackay. 2000. Quantitative trait loci for life span in *Drosophila melanogaster*: Interactions with genetic background and larval density. Genetics 155:1773–1788.

Lewis, Z., N. Wedell, and J. Hunt. 2011. Evidence for strong intralocus sexual conflict in the Indian meal moth, *Plodia interpunctella*. Evolution 65:2085–2097.

Mallet, M. A., and A. K. Chippindale. 2011. Inbreeding reveals stronger net selection on *Drosophila melanogaster* males: implications for mutation load and the fitness of sexual females. Heredity 106:994–1002.

Martinossi-Allibert, I., G. Arnqvist, and D. Berger. 2017. Sex-specific selection under environmental stress in seed beetles. Journal of Evolutionary Biology 30:161–173.

Martinossi-Allibert, I., U. Savkovic, M. Dordevic, G. Arnqvist, B. Stojkovic, and D. Berger. 2018. The consequences of sexual selection in well-adapted and maladapted populations of bean beetles. Evolution 72:518–530.

McCleery, R. H., R. A. Pettifor, P. Armbruster, K. Meyer, B. C. Sheldon, and C. M. Perrins. 2004. Components of variance underlying fitness in a natural population of the great tit *Parus major*. American Naturalist 164:E62–E72.

McFarlane, S. E., J. C. Gorrell, D. W. Coltman, M. M. Humphries, S. Boutin, and A. G. McAdam. 2014. Very low levels of direct additive genetic variance in fitness and fitness components in a red squirrel population. Ecology and Evolution 4:1729–1738.

Merila, J., and B. C. Sheldon. 2000. Lifetime reproductive success and heritability in nature. American Naturalist 155:301–310.

Moorad, J. A., and C. A. Walling. 2017. Measuring selection for genes that promote long life in a historical human population. Nature Ecology & Evolution 1:1773–1781.

Muhlhauser, C., and W. U. Blanckenhorn. 2004. The quantitative genetics of sexual selection in the dung fly *Sepsis cynipsea*. Behaviour 141:327–341.

Pélissié, B., P. Jarne, and P. David. 2012. Sexual selection without sexual dimorphism: Bateman gradients in a simultaneous hermaphrodite. Evolution 66:66–81.

Pettay, J. E., L. E. B. Kruuk, J. Jokela, and V. Lummaa. 2005. Heritability and genetic constraints of life-history trait evolution in preindustrial humans. Proceedings of the National Academy of Sciences of the United States of America 102:2838–2843.

Poissant, J., M. B. Morrissey, A. G. Gosler, J. Slate, and B. C. Sheldon. 2016. Multivariate selection and intersexual genetic constraints in a wild bird population. Journal of Evolutionary Biology 29:2022–2035.

Punzalan, D., M. Delcourt, and H. D. Rundle. 2014. Comparing the intersex genetic correlation for fitness across novel environments in the fruit fly, *Drosophila serrata*. Heredity 112:143–148.

Qvarnstrom, A., J. E. Brommer, and L. Gustafsson. 2006. Testing the genetics underlying the co-evolution of mate choice and ornament in the wild. Nature 441:84–86.

Rapkin, J., C. R. Archer, C. E. Grant, K. Jensen, C. M. House, A. J. Wilson, and J. Hunt. 2017. Little evidence for intralocus sexual conflict over the optimal intake of nutrients for life span and reproduction in the black field cricket *Teleogryllus commodus*. Evolution 71:2159–2177.

Rodriguez-Munoz, R., A. Bretman, J. D. Hadfield, and T. Tregenza. 2008. Sexual selection in the cricket *Gryllus bimaculatus*: no good genes? Genetica 134:129–136.

Ruzicka, F., M. S. Hill, T. M. Pennell, I. Flis, F. C. Ingleby, R. Mott, K. Fowler et al. 2019. Genome-wide sexually antagonistic variants reveal long-standing constraints on sexual dimorphism in fruit flies. Plos Biology 17.

Tarka, M., M. Akesson, D. Hasselquist, and B. Hansson. 2014. Intralocus sexual conflict over wing length in a wild migratory bird. American Naturalist 183:62–73.

Teplitsky, C., J. A. Mills, J. W. Yarrall, and J. Merila. 2009. Heritability of fitness components in a wild bird population. Evolution 63:716–726.

Vermeulen, C. J., R. Bijlsma, and V. Loeschcke. 2008. A major QTL affects temperature sensitive adult lethality and inbreeding depression in life span in *Drosophila melanogaster*. Bmc Evolutionary Biology 8.

Vieira, C., E. G. Pasyukova, Z. B. Zeng, J. B. Hackett, R. F. Lyman, and T. F. C. Mackay. 2000. Genotype-environment interaction for quantitative trait loci affecting life span in *Drosophila melanogaster*. Genetics 154:213–227.

Walling, C. A., M. B. Morrissey, K. Foerster, T. H. Clutton-Brock, J. M. Pemberton, and L. E. B. Kruuk. 2014. A multivariate analysis of genetic constraints to life history evolution in a wild population of red deer. Genetics 198:1735-+.

Wayne, M. L., J. B. Hackett, C. L. Dilda, S. V. Nuzhdin, E. G. Pasyukova, and T. F. C. MacKay. 2001. Quantitative trait locus mapping of fitness-related traits in *Drosophila melanogaster*. Genetical Research 77:107–116.

Wheelwright, N. T., L. F. Keller, and E. Postma. 2014. The effect of trait type and strength of selection on heritability and evolvability in an island bird population. Evolution 68:3325–3336.

Wolak, M. E., P. Arcese, L. F. Keller, P. Nietlisbach, and J. M. Reid. 2018. Sex-specific additive genetic variances and correlations for fitness in a song sparrow (*Melospiza melodia*) population subject to natural immigration and inbreeding. Evolution 72:2057–2075.

Zajitschek, F., J. Hunt, S. R. K. Zajitschek, M. D. Jennions, and R. Brooks. 2007. No intra-locus sexual conflict over reproductive fitness or ageing in field crickets. Plos One 2.

Zietsch, B. P., R. Kuja-Halkola, H. Walum, and K. J. H. Verweij. 2014. Perfect genetic correlation between number of offspring and grandoffspring in an industrialized human population. Proceedings of the National Academy of Sciences of the United States of America 111:1032–1036.

